# The light chain of the L9 antibody is critical for binding circumsporozoite protein minor repeats and preventing malaria

**DOI:** 10.1101/2021.08.11.455919

**Authors:** Lawrence T. Wang, Nicholas K. Hurlburt, Arne Schön, Barbara J. Flynn, Lais S. Pereira, Marlon Dillon, Yevel Flores-Garcia, Brian Bonilla, Fidel Zavala, Azza H. Idris, Joseph R. Francica, Marie Pancera, Robert A. Seder

**Author notes:** These authors contributed equally (Lawrence T. Wang, Nicholas K. Hurlburt, Marie Pancera, Robert A. Seder). Correspondence (M.P.), (R.A.S.).

## Abstract

L9 is a potent human monoclonal antibody (mAb) that preferentially binds two adjacent NVDP minor repeats and cross-reacts with NANP major repeats of the *Plasmodium falciparum* circumsporozoite protein (PfCSP) on malaria-infective sporozoites. Understanding this mAb’s ontogeny and mechanisms of binding PfCSP to neutralize sporozoites will facilitate vaccine development. Here, we isolated mAbs clonally related to L9 and showed that this B-cell lineage has baseline NVDP affinity and evolves to acquire NANP reactivity. Pairing the L9 kappa light chain (L9κ) with clonally-related heavy chains resulted in chimeric mAbs that cross-linked two NVDP, cross-reacted with NANP, and more potently neutralized sporozoites compared to their original light chain. Structural analyses revealed that chimeric mAbs bound the minor repeat motif in a type-1 β-turn seen in other repeat-specific antibodies. These data highlight the importance of L9κ in binding NVDP on PfCSP to neutralize SPZ and suggest that PfCSP-based immunogens might be improved by presenting ≥2 NVDP.

## INTRODUCTION

Interventions like vaccines are needed to prevent malaria, a parasitic mosquito-borne disease that killed ∼409,000 people in 2019, mostly from *Plasmodium falciparum* (Pf) (WHO, 2020). The most advanced malaria vaccine is RTS,S, a protein subunit vaccine that prevents malaria by inducing antibodies against the Pf circumsporozoite protein (PfCSP), the dominant surface protein on infectious sporozoites (SPZ) spread by mosquitoes (Cockburn and Seder, 2018). RTS,S adjuvanted with AS01 confers ∼ 50% protection against clinical disease at one year (Olotu et al., 2013, 2016), which declines over time as anti-PfCSP antibodies wane (White et al., 2015). Thus, PfCSP-based immunogens must be improved to increase vaccine efficacy.

PfCSP is an optimal vaccine target since it is required for SPZ to infect hepatocytes (Cerami et al., 1992). PfCSP has three domains: an N-terminus, a central domain composed of repeating tetrapeptides, and a C-terminus (Bowman et al., 1999). In the Pf 3D7 reference strain, the region at the junction of the N-terminus and repeat domain commences with NPDP followed by 3 alternating NANP and NVDP repeats. This “junctional region” is followed by 35 NANP repeats, with a fourth NVDP inserted after the twentieth NANP (Cockburn and Seder, 2018). Structural studies indicate that the motifs recognized by PfCSP mAbs in the repeat domain are DPNA, NPNV, and NPNA created by the joining of the three tetrapeptides (Oyen et al., 2018; Plassmeyer et al., 2009). Notably, RTS,S contains only 19 NANP repeats and the C-terminus (Stoute et al., 1997).

The immunodominant NANP repeats are targeted by nearly all neutralizing PfCSP mAbs reported so far (Julien and Wardemann, 2019). However, the isolation of rare and potent mAbs that preferentially bind NPDP (Kisalu et al., 2018; Tan et al., 2018) or the NVDP repeats (Wang et al., 2020) identified these subdominant epitopes as new sites of vulnerability on PfCSP. As these epitopes are not contained in RTS,S, these discoveries led to the development of next-generation vaccines against the junctional region (Atcheson et al., 2021; Calvo-Calle et al., 2021; Francica et al., 2021; Jelínková et al., 2021).

We recently compared the binding and potency of a panel of protective human PfCSP mAbs to determine which epitopes might improve immunogen design and select the most potent mAb for clinical development. mAb L9, which preferentially binds NVDP repeats and cross-reacts with NANP repeats, was shown to be the most potently protective mAb in the panel. Notably, L9 and other potent mAbs bound recombinant PfCSP (rPfCSP) in two binding events with distinct affinities by isothermal titration calorimetry (ITC), suggesting that this *in vitro* signature of “two-step binding” may be correlated with *in vivo* SPZ neutralization (Wang et al., 2020).

Similar to the majority of human PfCSP repeat mAbs, L9 is encoded by immunoglobulin (Ig) V_H_3-33 and Vκ1-5 (heavy and kappa chain variable domains) (Julien and Wardemann, 2019). Most V_H_3-33/Vκ1-5 mAbs preferentially bind NANP repeats and express an 8–amino-acid–long Vκ complementarity-determining region 3 (KCDR3:8) and a tryptophan at position 52 in CDRH2 (H.W52) (Imkeller et al., 2018; Murugan et al., 2018, 2020). Furthermore, these NANP-preferring mAbs require a minimal epitope of three NANP (Oyen et al., 2017) and rPfCSP binding is abrogated when V_H_3-33 is paired with Vκ1-5 with KCDR3:9 (Imkeller et al., 2018). Conversely, L9 expresses KCDR3:9 but binds rPfCSP with high affinity, preferentially binds NVDP repeats, and can only bind NANP repeats if they are sufficiently concatenated (Wang et al., 2020). These data suggest that V_H_3-33/Vκ1-5 mAbs can also target NVDP repeats; however, it is unclear whether the L9 B-cell lineage originated as a NANP binder that mutated to acquire NVDP affinity, as has been suggested for NPDP-reactive mAbs (Tan et al., 2018, 2019), or vice versa.

Here, to elucidate how the L9 B-cell lineage developed, mAbs clonally related to L9 were isolated. These mAbs preferentially bound NVDP repeats and had minimal-to-undetectable NANP reactivity. Pairing the L9 Igκ light chain (L9κ) with clonally-related Ig_H_, compared to their original Igκ, resulted in chimeric mAbs with similar binding properties to L9 (i.e., binding two adjacent NVDP repeats, cross-reacting with NANP, and two-step binding rPfCSP) and improved SPZ neutralization *in vivo*. Structures of antigen-binding fragments (Fabs) from two chimeric mAbs in complex with a minimal peptide NANPNVDP showed nearly identical binding mechanisms, with the NPNV adopting a type-1 β-turn. This study provides insight into the evolution of antibodies targeting the subdominant NVDP minor repeats and the binding properties associated with the potent SPZ neutralization of L9.

## RESULTS

### Isolation of a panel of mAbs clonally related to L9

L9 was isolated from a volunteer immunized with radiation-attenuated PfSPZ (Lyke et al., 2017; Wang et al., 2020). To isolate clonally-related mAbs, CD3^-^CD20^-^CD19^+^CD27^+^CD38^+^ plasmablasts were sorted from the same volunteer 7 days after the second and third immunizations. A total of 480 plasmablasts were isolated, from which 194 high-quality IgG V_H_ sequences were cloned. These sequences were compared to the L9 V_H_ (L9_H_) to identify clonally-related sequences. Two V_H_ (D2_H_ and F10_H_) from plasmablasts collected after the second immunization were clonally related to L9_H_. Specifically, all three V_H_ were encoded by V_H_3-33*01/06, J_H_4*02, and had a 13-amino-acid-long HCDR3; furthermore, the HCDR3 of D2_H_ and F10_H_ were 85% identical to L9_H_ (Table S1).

**Table 1.** ITC of NVDP-preferring mAbs binding to peptides or rPfCSP. . ITC measurements of the stoichiometry (N), dissociation constant (K_D_), change in Gibbs free energy (ΔG), change in enthalpy (ΔH), and change in entropy contribution to Gibbs free energy (−TΔS) of indicated mAb (Fab or IgG) binding to peptides (pep 22, NA<u>NPNV</u>DPNA<u>NPNV</u>D; pep 25, NVDPNA<u>NPNV</u>DPNAN; pep 29, NANPNANPNANPNAN; minimal peptide, NA<u>NPNV</u>DP) and rPfCSP constructs (FL, 1 NPDP, 4 NVDP, 38 NANP; 5/3, 1 NPDP, 3 NVDP, 5 NANP; Δ(NVDP)_4_, 1 NPDP, 42 NANP). “-” indicate undetectable parameters. Tabulated values were averaged from 2-3 independent experiments. L9 and 317 IgG data were taken from (Wang et al., 2020).

| mAb | Antigen | Binding Event 1 |  |  |  |  | Binding Event 2 |  |  |  |  |
| --- | --- | --- | --- | --- | --- | --- | --- | --- | --- | --- | --- |
|  |  | N<sub>1</sub> | K<sub>D1</sub><br>(nM) | ΔG<sub>1</sub><br>(kcal/mol) | ΔH<sub>1</sub><br>(kcal/mol) | -TΔS<sub>1</sub><br>(kcal/mol) | N<sub>2</sub> | K<sub>D2</sub><br>(nM) | ΔG<sub>2</sub><br>(kcal/mol) | ΔH<sub>2</sub><br>(kcal/mol) | -TΔS<sub>2</sub><br>(kcal/mol) |
| L9 Fab | Pep 22 | 1.6 | 13 | -10.8 | -15.8 | +5.0 | - | - | - | - | - |
|  | Pep 25 | 1.0 | 1000 | -8.2 | -13.8 | +5.6 | - | - | - | - | - |
|  | Pep 29 | - | - | - | - | - | - | - | - | - | - |
|  | NANPNVDP | 0.7 | 1900 | -7.8 | -6.4 | -1.4 | - | - | - | - | - |
| L9 IgG | FL | 1.9 | 22 | -10.4 | -30.0 | +19.6 | 9.3 | 47 | -10.0 | -9.1 | -0.9 |
|  | 5/3 | 3.1 | 3 | -11.6 | -17.0 | +5.4 | - | - | - | - | - |
|  | Δ(NVDP)<sub>4</sub> | 12.7 | 73 | -9.7 | -12.4 | +2.7 | - | - | - | - | - |
| L9<sub>H</sub>F10<sub>k</sub> IgG | FL | 2.5 | 58 | -9.9 | -11.7 | +1.8 | - | - | - | - | - |
|  | 5/3 | 2.2 | 2.1 | -11.8 | -18.2 | +6.4 | - | - | - | - | - |
|  | Δ(NVDP)<sub>4</sub> | - | - | - | - | - | - | - | - | - | - |
| F10<sub>H</sub>L9<sub>k</sub> IgG | FL | 3.4 | 7 | -11.1 | -14.1 | +3.0 | 6.8 | 400 | -8.7 | -10.3 | +1.6 |
|  | 5/3 | 2.4 | 1.2 | -12.2 | -15.8 | +3.6 | - | - | - | - | - |
|  | Δ(NVDP)<sub>4</sub> | 8.1 | 96 | -9.6 | -9.9 | +0.3 | - | - | - | - | - |
| F10 IgG | FL | 3.4 | 14 | -10.7 | -13.4 | +2.7 | - | - | - | - | - |
|  | 5/3 | 2.1 | 4 | -11.4 | -17.5 | +6.1 | - | - | - | - | - |
|  | Δ(NVDP)<sub>4</sub> | - | - | - | - | - | - | - | - | - | - |
| D2<sub>H</sub>L9<sub>k</sub> IgG | FL | 3.6 | 5 | -11.3 | -9.2 | -2.1 | 7.7 | 270 | -9.0 | -6.5 | -2.5 |
|  | 5/3 | 2.2 | 1.6 | -12.0 | -13.0 | +1.0 | - | - | - | - | - |
|  | Δ(NVDP)<sub>4</sub> | 9.7 | 87 | -9.6 | -7.3 | -2.3 | - | - | - | - | - |
| D2<sub>H</sub>F10<sub>k</sub> IgG | FL | 2.7 | 32 | -10.2 | -11.5 | +1.3 | - | - | - | - | - |
|  | 5/3 | 2.3 | 6.5 | -11.2 | -15.6 | -4.4 | - | - | - | - | - |
|  | Δ(NVDP)<sub>4</sub> | - | - | - | - | - | - | - | - | - | - |
| 317 IgG | FL | 7.5 | 2 | -11.9 | -26.3 | +14.4 | 4.7 | 17 | -10.6 | -16.7 | +6.1 |
|  | 5/3 | 0.5 | 2 | -11.9 | -28.0 | +16.1 | 1.2 | 7 | -11.1 | -16.7 | +5.6 |
|  | Δ(NVDP)<sub>4</sub> | 8.5 | 2 | -11.8 | -23.3 | +11.5 | 4.1 | 14 | -10.7 | -17.6 | +6.9 |

Regarding light chains, only F10κ was cloned. F10κ and L9κ were Vκ1-5*03, Jκ1*01, and had a 9-amino-acid-long KCDR3; furthermore, the KCDR3 of F10κ was 89% identical to L9κ (Table S1). The V_H_/Vκ sequences of D2, F10, and L9 were used to infer their most recent common ancestor (L9_MRCA_) V_H_/Vκ, which were respectively 0.35% and 0% diverged from germline. Notably, the V_H_/Vκ of L9 were more somatically mutated than those of D2 and F10 (Table S1) and all V_H_ genes encoded tryptophan at position 52 (H.W52) in HCDR2, which has been shown to be critical for binding NANP repeats (Imkeller et al., 2018). To elucidate the contribution of Igκ to mAb recognition of PfCSP and resultant SPZ neutralization, chimeric mAbs were created wherein L9_H_ was paired with F10κ (L9_H_F10κ) and F10_H_ was paired with L9κ (F10_H_L9κ). As D2κ was not recovered, L9κ and F10κ were paired with D2_H_ to respectively create the chimeric mAbs D2_H_L9κ and D2_H_F10κ (Figure 1A).

**Figure 1.**
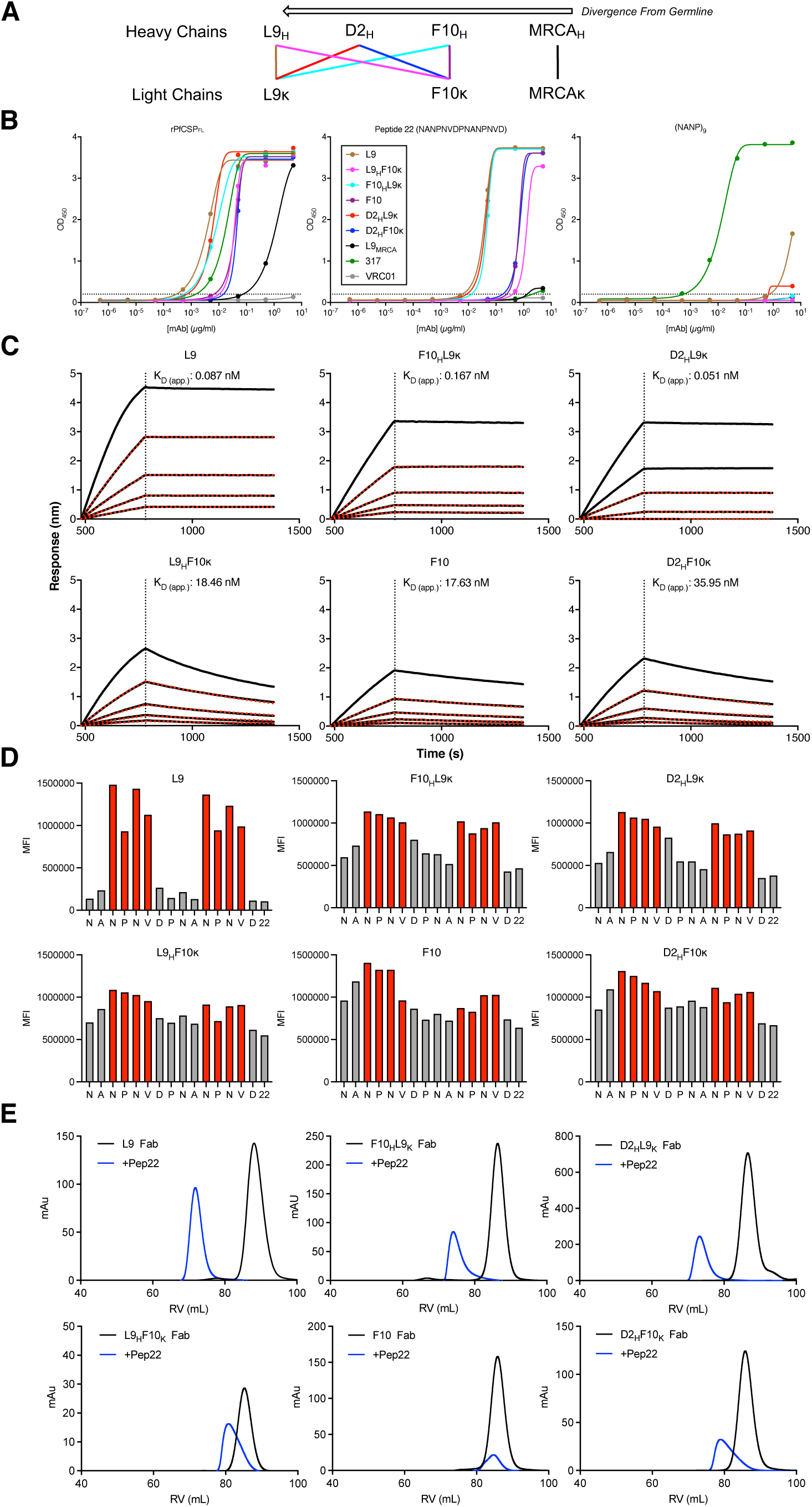
L9κ enables binding to two adjacent NPNV minor repeat motifs. (A) Ig heavy and light chain pairings matched to create normal mAbs (L9, F10, L9_MRCA_) and chimeric mAbs (L9_H_F10κ, F10_H_L9κ, D2_H_L9κ, D2_H_F10κ) used in this study. Each heavy and light chain’s divergence from germline is depicted. (B) Binding of various concentrations of indicated mAbs to rPfCSP_FL_ (left), peptide 22 (NANPNVDPNANPNVD, middle), and (NANP)_9_ (right) measured by ELISA; optical density at 450 nm (OD_450_) is plotted. The NANP-preferring PfCSP mAb 317 and the anti-gp120 mAb VRC01 were included as positive and negative controls, respectively. (C) Binding response (nm) and apparent avidity (KD_app_, nM) of indicated mAbs binding to peptide 22 determined through BLI. (D) Competition ELISA of indicated mAbs binding to rPfCSP_FL_ and various concentrations of peptide 22 (rightmost bar) or variant peptides (other bars) where the indicated amino acid was mutated to alanine or serine. Area under the curve (AUC) is plotted. (E) SEC plots of indicated Fabs incubated with peptide 22 (Fabs alone, black peak; Fabs+pep22, blue peak). All data are representative of two independent experiments.

### The L9 B-cell lineage preferentially binds NPNV minor repeat motifs

All mAbs in the panel were assessed for their binding to full-length rPfCSP (rPfCSP_FL_); the NANP-preferring PfCSP mAb 317 (Oyen et al., 2017) and the anti-HIV-1 mAb VRC01 (Zhou et al., 2010) were respectively included as positive and negative controls. All mAbs bound rPfCSP_FL_, confirming that mismatching Igκ with Ig_H_ did not interfere with antigen recognition by the chimeric mAbs (Figure 1B). Notably, L9κ-containing mAbs (L9, F10_H_L9κ, D2_H_L9κ) had higher rPfCSP_FL_ binding compared to F10κ-containing mAbs (F10, L9_H_F10κ, D2_H_F10κ).

L9_MRCA_ had the lowest rPfCSP_FL_ binding, likely due to its low divergence from germline (Table S1).

L9 was reported to preferentially bind the two adjacent NPNV motifs associated with the NVDP repeats in the 15mer peptide 22 (NA<u>NPNV</u>DPNA<u>NPNV</u>D) and weakly cross-react with the eight concatenated NPNA motifs in the 36mer peptide (NANP)_9_ (Wang et al., 2020). To determine if the mAbs in the panel had similar binding properties, we measured their peptide 22 and (NANP)_9_ binding by ELISA (Figure 1B). Similar to L9, all mAbs demonstrated higher binding to peptide 22 than to (NANP)_9_. As with rPfCSP_FL_, L9κ-containing mAbs had higher peptide 22 binding than F10κ-containing mAbs and L9_MRCA_ had lower, but detectable, peptide 22 binding. The only mAbs with weak, though detectable, (NANP)_9_ binding were L9 and D2_H_L9κ.

To quantify the mAb panel’s avidity for peptide 22, biolayer interferometry (BLI) was used to measure their apparent avidity (K_Dapp_) for peptide 22 (Figure 1C). Consistent with the ELISA data, L9κ-containing mAbs had K_Dapp_ 100-700-fold lower than F10κ-containing mAbs, suggesting that L9κ improves mAb avidity for peptide 22. Together, these data show that the mAbs in the panel preferentially bind the NVDP-containing peptide 22 over the NANP- containing (NANP)_9_ and that, compared to F10κ, L9κ improves binding to rPfCSP_FL_, peptide 22, and (NANP)_9_.

To further characterize the mAb panel’s interactions with peptide 22, alanine scanning mutagenesis was used to define the critical residues within the peptide 22 sequence bound by each mAb (Figure 1D). Unlike L9 IgG, which clearly bound both NPNV motifs, F10 IgG mostly bound the first NPNV motif in peptide 22 and did not appear to bind the second NPNV. Remarkably, L9κ-containing IgG clearly bound both NPNV motifs while F10κ-containing IgG had more equivocal NPNV binding, with binding being targeted more towards the first NPNV. Furthermore, size exclusion chromatography (SEC) of Fabs bound to peptide 22 showed that L9κ-containing Fab-peptide 22 complexes eluted earlier (i.e., were larger) than F10κ-containing Fab-peptide 22 complexes, suggesting that peptide 22 is bound by two L9κ-containing Fabs but only one F10κ-containing Fab (Figure 1E). These data show that the higher peptide 22 avidity of L9κ-containing mAbs compared to F10κ-containing mAbs is because L9κ enables high-affinity binding to two adjacent NPNV motifs instead of one.

To extend the analysis for L9, ITC was used to measure the affinity and stoichiometry of L9 Fabs binding to 15mer peptides 22, 25 (NVDPNA<u>NPNV</u>DPNAN), and 29 (NANPNANPNANPNAN), as well as a minimal 8mer peptide (NA<u>NPNV</u>DP) (Table 1). L9 bound ∼2 binding sites with an affinity of 13 nM on peptide 22 (two NPNV) and ∼1 binding site with an affinity of 1,000 and 1,900 nM on peptide 25 and NANPNVDP, respectively (one NPNV). L9 had no detectable binding to peptide 29, which contains 3 NPNA motifs. Collectively, these data confirm that L9 lineage mAbs recognize NPNV instead of NPNA, that L9 IgG requires two adjacent NPNV motifs for high-affinity binding to NVDP-containing peptides, and that pairing L9κ with closely-related Ig_H_ is sufficient to enable cross-linking of two NPNV motifs.

### F10_H_L9κ and L9_H_F10κ Fabs bind NPNV motifs in an identical manner

Next, we sought to structurally determine the binding mechanism of L9 using X-ray crystallography, but diffracting crystals could only be generated in the absence of peptides and a structure of L9 Fab alone (apo-L9) was solved to 2.93 Å (Figure S1A). To gain structural insight into the NPNV binding mechanism of L9, we crystallized Fabs of F10_H_L9κ and L9_H_F10κ in complex with the NANPNVDP peptide and solved their structures to 1.89 Å and 2.23 Å, respectively (Figure 2A,C). F10_H_L9κ Fab binds NANPNVDP with the NPNV motif adopting a type-1 β-turn (Figure S1B) that the NPNA motif was shown to adopt (Ghasparian et al., 2006) and is commonly found in NPNA-preferring repeat mAbs (Figure 2B) (Imkeller et al., 2018; Oyen et al., 2017; Pholcharee et al., 2021). All three HCDRs and the KCDR1 and KCDR3 bind NANPNVDP with a total buried surface area (BSA) of ∼373 Å^2^, ∼203 Å^2^ from the Ig_H_ and ∼170 Å^2^ from Igκ (Figure 2E). The two Asn residues form a network of hydrogen bonds with both Ig_H_/Igκ, with the Val sitting in a hydrophobic pocket formed by three aromatic residues on the KCDR1, KCDR3, and HCDR3 (Figure 2B). Surprisingly, the structure of L9_H_F10κ Fab- NANPNVDP showed an almost identical binding mechanism to F10_H_L9κ Fab-NANPNVDP (Figure 2D,F). The NPNV adopts the type-1 β-turn structure and is bound with a total BSA of ∼364 Å^2^, ∼196 Å^2^ from Ig_H_ and ∼168 Å^2^ from Igκ (Figure 2E).

**Figure 2.**
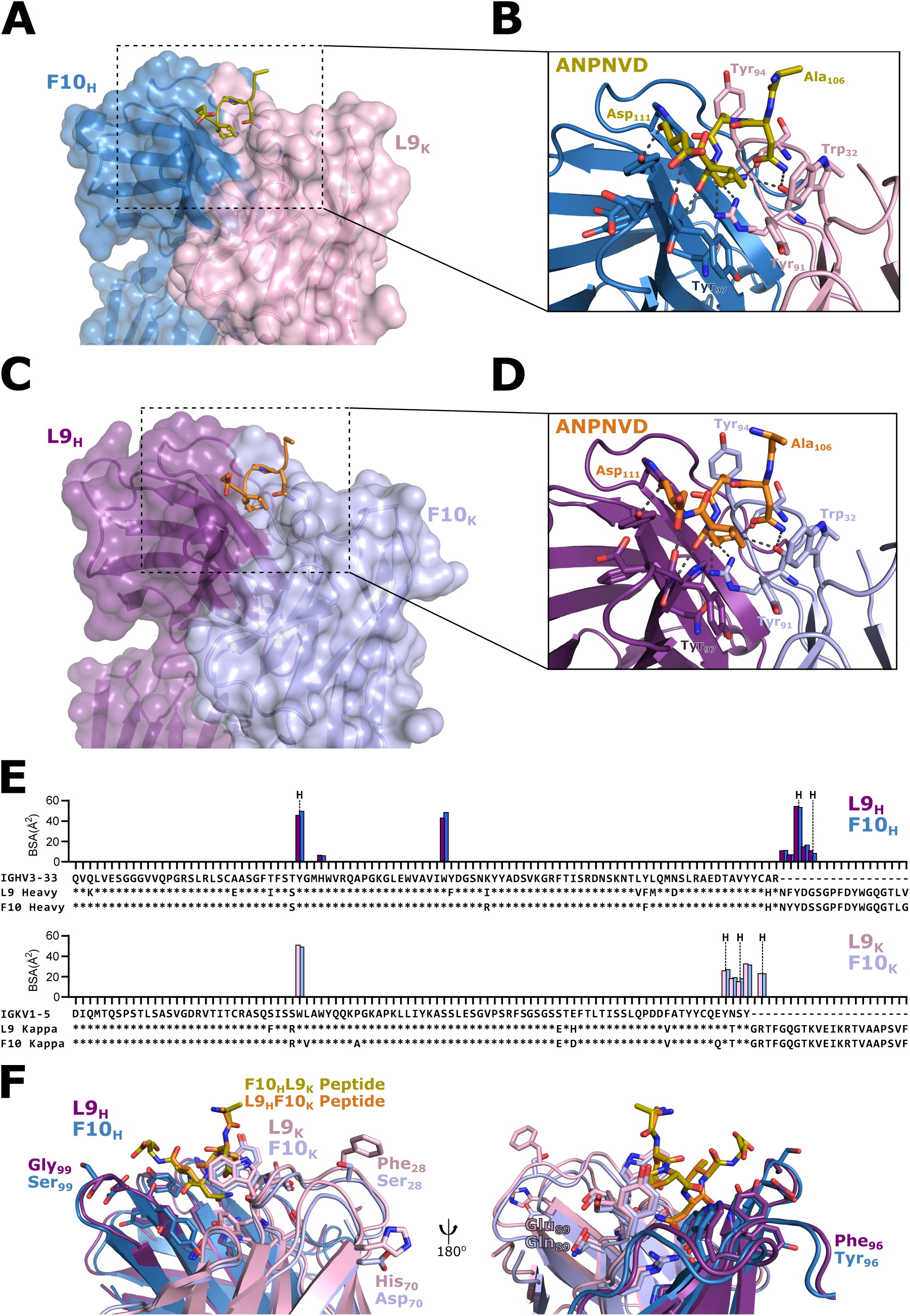
Crystal structures of F10_H_L9κ and L9_H_F10κ in complex with the NANPNVDP peptide. (A) Structures of F10_H_L9κ Fab and (C) L9_H_F10κ Fab bound to NANPNVDP. The structures are shown in a cartoon representation with a transparent surface representation. The sidechains of the peptides are shown. (B) and (D) respectively show zoomed-in images of the binding sites in A and C. Peptide interacting residues are shown in stick representations, with hydrogen bonds represented by dashed lines. Named residues are depicted to orient structure to location of CDRs. (E) BSA plots of each Fab residue interacting with the peptide and a sequence alignment with the germline V-genes. Residues involved in hydrogen bonding are marked with a “H”. (F) Structural alignment of the F10_H_L9κ and L9_H_F10κ Fab structures using the NANPNVDP peptide. Residues that differ between the two structures are labeled.

To further investigate the differences between L9_H_F10κ and F10_H_L9κ, we aligned both structures by their shared cognate NANPNVDP peptide. The two peptides align with a root mean square deviation (RMSD) of 0.189 Å^2^ over six Cα atoms (Figure 2F). The only two residues that differ in the binding site, positions 96 (L9:Phe and F10:Tyr) and 99 (L9:Gly and F10:Ser) in Ig_H_ by kabat numbering, are both main chain interactions and are unlikely to contribute to differences between the two chimeric mAbs. The interacting residues in both Igκ are identical.

There are three differences in Igκ that may contribute to the increased binding potential of L9κ. In the KCDR3, Glu_90_ in L9κ is charge swapped to a Gln in F10κ and is the only difference near the binding site. Two other differences occur in KCDR1 and framework region three where Ser_28_ and Asp_70_ in F10κ are mutated to aromatic residues in L9κ, Phe_28_ and His_70_, both of which are solvent exposed. All three of these residues in L9κ are unusual mutations, occurring in less than 1% of kappa chain sequences (abYsis). These structures likely represent the binding mechanism of L9 to NPNV, but further structural information is needed to determine why L9κ enables binding to two adjacent NPNV motifs, as in peptide 22.

### The L9 B-cell lineage has baseline NPNV affinity

To further define the evolution of epitope reactivities in mAbs from the L9 B-cell lineage, the binding of L9_MRCA_, F10, and L9 to three rPfCSP constructs (FL, 5/3, Δ(NVDP)_4_; Figure S2A) were measured by ELISA; 317 was included as a NANP-preferring control mAb. rPfCSP_5/3_ is a truncated construct containing only the junctional region (i.e., NPDP followed by 5 NANP interspersed with 3 NVDP repeats); rPfCSP_Δ(NVDP)4_ has 1 NPDP and 42 NANP repeats, as all NVDP were changed to NANP.

F10 and L9 had high rPfCSP_FL_ and rPfCSP_5/3_ binding but only L9 had detectable rPfCSP_Δ(NVDP)4_ binding (Figure S2B). Notably, L9_MRCA_ only bound rPfCSP_FL_ and rPfCSP_5/3_ and had no measurable rPfCSP_Δ(NVDP)4_ binding. Given these three mAbs’ relative degrees of somatic mutation (L9 > F10 > L9_MRCA_, with L9_MRCA_ being near-germline; Table S1), these data suggest that the L9 lineage has baseline affinity for the junctional region and mutates to gain NANP cross-reactivity.

The binding preferences of the L9 lineage were corroborated by competition ELISA showing that a junctional peptide more potently competed mAb binding to rPfCSP_FL_ and rPfCSP_5/3_ than (NANP)_9_, with the degree of competition correlating to the mAbs’ somatic mutations (Figure S2C). Notably, L9 was the only mAb that detectably bound rPfCSP_Δ(NVDP)4_ and the junctional peptide more potently competed L9 binding than (NANP)_9_.

To extend the epitope mapping analysis, peptides 20-61 spanning the repeat domain were used to compete L9_MRCA_, F10, and L9 binding to rPfCSP_FL_ (Figure S2D). These data confirmed the markedly high affinity of L9 for peptide 22, the relatively high binding of F10 and L9_MRCA_ to peptides 21/22/23/43 (all contain ≥1 NPNV), and all three mAbs’ low/undetectable binding to peptides 27/29/61 (all lack NPNV). Furthermore, all three mAbs preferentially bound (NPNV)_4_ compared to (NPNA)_4_, confirming that the core motif recognized by these mAbs is NPNV. Collectively, these data show that the L9 V_H_3-33/Vκ1-5 B-cell lineage originated as subdominant NPNV binders that somatically mutated to acquire affinity for immunodominant NPNA motifs.

### L9κ confers NPNA cross-reactivity and two-step binding to rPfCSP

To elucidate how the mAb panel’s relative affinities for NPNV and NPNA motifs impacted their binding to full-length protein, ITC was used to measure their binding stoichiometry and affinity to rPfCSP_FL_, rPfCSP_5/3_, and rPfCSP_Δ(NVDP)4_. We previously reported that the most potent mAbs for protecting against *in vivo* SPZ challenge, including L9, bound rPfCSP_FL_ in two binding events with distinct affinities (termed “two-step binding”) and displayed high-affinity binding to the junctional region in rPfCSP_5/3_. Furthermore, we showed that L9 binds rPfCSP_Δ(NVDP)4_ in a single step with lower affinity, confirming that L9 requires NPNV to two-step bind rPfCSP but can cross-react with NPNA motifs (Wang et al., 2020).

Remarkably, L9κ-containing mAbs (L9, F10_H_L9κ, D2_H_L9κ) bound rPfCSP_FL_ in two steps while F10κ-containing mAbs (F10, L9_H_F10κ, D2_H_F10κ) bound in a single step, resulting in the higher stoichiometry of L9κ-containing mAbs for rPfCSP_FL_ (Table 1). The mAbs’ stoichiometry in binding event 1 (N_1_) for rPfCSP_FL_ and rPfCSP_5/3_ were similar (∼2-3 binding sites), suggesting that the epitopes bound by these mAbs in binding event 1 are contained in rPfCSP_5/3_ (Table 1). Furthermore, only L9κ-containing mAbs had detectable binding to rPfCSP_Δ(NVDP)4_ with L9 having the highest stoichiometry (∼13 binding sites).

The mAb panel’s affinity in binding event 1 (K_D1_) was better for rPfCSP_5/3_ (1.2-6.5 nM) compared to rPfCSP_FL_ (5-58 nM), further indicating that binding event 1 is directed toward rPfCSP_5/3_ (Table 1). Consistent with the (NANP)_9_ ELISA data (Figure 1B), only L9κ-containing mAbs had detectable affinity for rPfCSP_Δ(NVDP)4_ with L9 having the best affinity, though significantly less than the NANP-preferring mAb 317 (73 vs. 2 nM; Table 1). Collectively, the ITC data confirm that all mAbs in the panel have high-affinity binding to NPNV motifs in the junctional region (i.e., rPfCSP_5/3_), that L9κ imparts lower-affinity binding to concatenated NPNA motifs (i.e., rPfCSP_Δ(NVDP)4_), and that two-step binding to rPfCSP_FL_ seen in L9κ-containing mAbs is due to their respective high- and low-affinity binding to NPNV and NPNA motifs.

### L9κ improves SPZ binding and neutralization

To determine whether the mAb panel’s stoichiometry and affinity for rPfCSP_FL_ correlate with their recognition of native PfCSP_FL_ on the surface of SPZ, flow cytometry was used to quantify mAb binding to transgenic *P. berghei* SPZ expressing PfCSP_FL_ and a green fluorescent protein/luciferase fusion protein (Pb-PfCSP-GFP/Luc-SPZ; hereafter Pb-PfCSP-SPZ). All mAbs in the panel bound Pb-PfCSP-SPZ (Figure 3A). Consistent with the rPfCSP_FL_ ELISA and ITC data (Figures 1A, Table 1), L9κ-containing mAbs (L9, F10_H_L9κ, D2_H_L9κ) had higher Pb-PfCSP-SPZ binding compared to their counterpart F10κ-containing mAbs (F10, L9_H_F10κ, D2_H_F10κ) while L9_MRCA_ had the lowest rPfCSP_FL_ binding.

**Figure 3.**
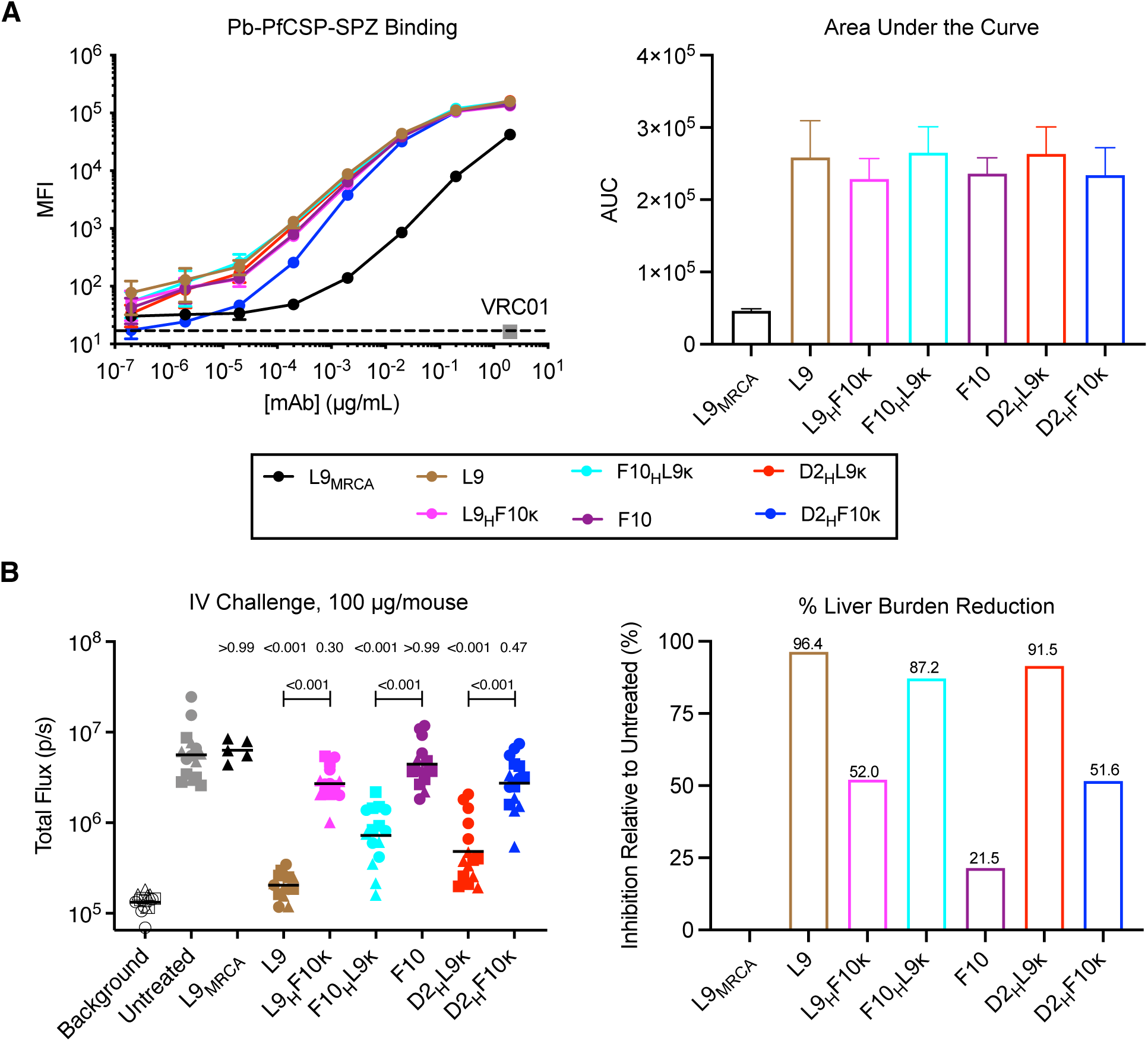
L9κ improves SPZ binding and neutralization. (A) Left: binding of various concentrations of indicated mAbs to Pb-PfCSP-SPZ determined by flow cytometry; median fluorescence intensity (MFI) of mAb-bound SPZ is plotted. Mean and standard error of the mean (SEM) MFI values from four independent experiments are shown except for L9_MRCA_, which was from two experiments. Right: area under the curve (AUC) and SEM values calculated from the MFI plots. (B) Left: reduction in parasite bioluminescence (total flux, photons/sec) in the livers of mice (n=15/group; data pooled from three independent experiments indicated by squares, circles, and triangles) mediated by 100 µg of indicated mAbs administered 2 hours before IV challenge with 2,000 Pb-PfCSP-SPZ. Top *P-*values were determined by comparing mAbs to untreated control using the Kruskal-Wallis test with Dunn’s post-hoc correction; bottom *P-* values reflect comparisons between L9κ-containing mAbs and their F10κ-containing counterparts using the two-tailed Mann–Whitney test. Right: percentage of liver burden reduction relative to untreated control mice mediated by each mAb.

We next evaluated the ability of each mAb in the panel to lower parasite liver burden in normal C57/BL6 mice challenged intravenously (IV) with Pb-PfCSP-SPZ, a model used to quantify SPZ neutralization by PfCSP mAbs *in vivo* (Raghunandan et al., 2020). Passive transfer of 100 µg L9κ-containing mAbs mediated 87-96% liver burden reduction relative to untreated controls, which was significantly greater than the 21-52% reduction provided by F10κ- containing mAbs (Figure 3B). Collectively, these data show that L9κ improves the ability of NVDP-preferring mAbs to bind and neutralize Pb-PfCSP-SPZ *in vivo*.

## DISCUSSION

This study showed that the Igκ of L9, a highly potent human PfCSP mAb that preferentially binds the subdominant NVDP minor repeats (Wang et al., 2020), is critical for this antibody’s unique binding properties and potent neutralization of SPZ. This finding was shown by pairing mAbs clonally related to L9 with different Ig_H_/Igκ to create L9κ-containing chimeric mAbs that recapitulated the binding properties of L9 (i.e., binds two adjacent NPNV motifs, cross-reacts with concatenated NPNA motifs, and two-step binds rPfCSP_FL_). Conversely, pairing L9_H_ and other V_H_ with the Igκ of a closely related mAb, F10κ, resulted in chimeric mAbs that bound only one NPNV motif and lacked NPNA affinity and two-step binding. Furthermore, these data confirm that two-step binding is detected by ITC when a mAb exhibits two distinct affinities for the junctional region and NANP repeats on rPfCSP_FL_, as has been previously suggested (Wang et al., 2020).

These data also show that cross-linking two adjacent NVDP repeats, cross-reacting with NANP repeats, and two-step binding rPfCSP are binding properties correlated with higher binding to native PfCSP on Pb-PfCSP-SPZ *in vitro* and improved protection against *in vivo* Pb-PfCSP-SPZ challenge. Importantly, F10κ-containing mAbs had high (1-7 nM) rPfCSP_5/3_ affinity comparable to L9κ-containing mAbs by ITC, suggesting that high-affinity binding to the junctional region alone may not be sufficient for potent SPZ neutralization. Further studies with transgenic SPZ lacking NVDP repeats are required to determine whether protection mediated by L9κ-containing mAbs is due to their high-affinity divalent binding of adjacent NVDP repeats or lower-affinity multivalent interactions with NANP repeats. Collectively, this study identifies discrete binding properties associated with SPZ neutralization by NVDP-preferring mAbs and suggest that future PfCSP-based vaccines should include ≥2 adjacent NVDP repeats (NA<u>NPNV</u>DPNAN<u>PNV</u>DP) to induce neutralizing minor repeat antibodies.

In terms of elucidating the evolution of human minor repeat antibodies, this study showed that the L9 V_H_3-33/Vκ1-5 B-cell lineage has baseline affinity for NVDP repeats in the junctional region and somatically mutated to acquire NANP cross-reactivity. Specifically, the L9 lineage germline binds one NPNV and evolves to bind two adjacent NPNV motifs and concatenated NPNA motifs. This evolutionary pathway differs from a previous study showing that a large panel of V_H_3-33/Vκ1-5 PfCSP human mAbs had baseline NANP affinity centered on the conserved (N/D)PNANPN(A/V) core motif and evolved to acquire NPDP/NVDP cross-reactivity (Murugan et al., 2020). Having shown that the core motif bound by L9 lineage mAbs is NPNV, our study suggests that some rare V_H_3-33/Vκ1-5 lineages initially target subdominant NVDP repeats and evolve later to cross-react with immunodominant NANP repeats. These data may assist in designing NVDP-containing PfCSP immunogens to expand naïve B-cells expressing the germline receptor for L9.

Lastly, structural studies showed that F10_H_L9κ and L9_H_F10κ both bound NPNV in the minimal peptide NA<u>NPNV</u>DP in a type-1 β-turn that was near-identical to the NPNA type-1 β-turn commonly observed for NANP-preferring mAbs (Imkeller et al., 2018; Oyen et al., 2017; Pholcharee et al., 2021), suggesting that recognition of single NPNV motifs does not differ between these mAbs. The convergent conformations adopted by these subtly different tetrapeptides is in line with a previous study which also observed that cross-reactive PfCSP mAbs bind different repeat epitopes in near-identical conformations (Murugan et al., 2020). Further structural studies on longer peptides containing ≥2 NPNV motifs and rPfCSP_FL_ are needed to elucidate how L9κ contributes to binding adjacent NPNV motifs with high affinity and cross-reacting with NPNA motifs, as well as how these binding properties contribute to SPZ neutralization.

## ACKNOWLEDGMENTS

A.S. was supported by contract HHSN261200800001E from the National Cancer Institute, NIH. Y.F-G. and F.Z. thank the Bloomberg Philanthropies for their continued support. We thank B. Kim Lee Sim and Stephen L. Hoffman for providing the PfSPZ Vaccine, David Ambrozak for assistance with sorting plasmablasts, L. Stamatatos for use of laboratory space and equipment, and the J. B. Pendleton Charitable Trust for its generous support of Formulatrix robotic instruments. The Berkeley Center for Structural Biology is supported in part by the Howard Hughes Medical Institute. The Advanced Light Source is a Department of Energy Office of Science User Facility under Contract No. DE-AC02-05CH11231. The Pilatus detector on 5.0.1. was funded under NIH grant S10OD021832. The ALS-ENABLE beamlines are supported in part by the NIH, National Institute of General Medical Sciences, grant P30 GM124169. Results shown in this report are derived from work performed at Argonne National Laboratory (ANL), Structural Biology Center (SBC) at the Advanced Photon Source (APS), under U.S. Department of Energy, Office of Biological and Environmental Research contract DE-AC02-06CH11357.

## AUTHOR CONTRIBUTIONS

L.T.W., N.K.H., J.F., M.P., and R.A.S. conceived the project, designed experiments, and wrote the manuscript. L.T.W, N.K.H., A.S., B.J.F., L.S.P., A.H.I., M.D., and B.B. cloned antibodies and/or conducted experiments. Y.F-G. and F.Z. provided sporozoites for flow cytometry. All authors reviewed, edited and approved the paper.

## MATERIALS & METHODS

### Lead Contact

Further information and requests for resources and reagents should be directed to and will be fulfilled by the Lead Contact, Robert A. Seder.

### Materials Availability

All unique reagents generated in this study are available from the Lead Contact with a completed Materials Transfer Agreement.

### Data and Code Availability

The heavy and light chain gene sequences of anti-PfCSP human mAbs isolated in this study were deposited in GenBank (Accession Numbers MZ686952 – MZ686956). The structure of apo-L9, F10_H_L9_K_-NANPNVDP, and L9_H_F10_K_-NANPNVDP were deposited in the Protein Data Bank (PBD), with PDB ID of 7RQP, 7RQQ, and 7RQR, respectively.

### Human clinical specimens

Clinical specimens were derived from healthy, malaria-naive adults (18-45 years of age) in VRC 314 clinical trial (https://clinicaltrials.gov/; NCT02015091) after obtaining written informed consent. Briefly, VRC 314 was a multi-institution, phase 1, open-label, dose-escalation trial with controlled human malaria infection that was designed to assess the safety, immunogenicity, and protective efficacy of the Sanaria PfSPZ Vaccine administered by intravenous or intramuscular injection (Lyke et al., 2017). The Sanaria PfSPZ Vaccine is composed of radiation-attenuated, aseptic, purified, cryopreserved *Plasmodium falciparum* sporozoites derived from the NF54 strain (Seder et al., 2013).

### Mice

Female 6- to 8-weeks old B6(Cg)-Tyrc-2J/J albino mice were obtained from The Jackson Laboratory. All mouse research was performed according to National Institutes of Health (NIH) guidelines for use and care of live animals approved by the institutional animal care and use ethics committees of the Vaccine Research Center (Animal Study Protocol VRC-17-702).

### Cell Lines

Expi293 and 293F cells used were from Thermo Fisher Scientific.

### Sporozoites

Transgenic *P. berghei* (strain ANKA 676m1c11, MRA-868) expressing full-length *P. falciparum* CSP and a green fluorescent protein/luciferase fusion protein (Pb-PfCSP-GFP/Luc-SPZ or Pb-PfCSP-SPZ) were obtained from salivary glands of infected mosquitoes, as previously described (Flores-Garcia et al., 2019). Briefly, *Anopheles stephensi* mosquitoes were fed on parasite-infected mice after confirming the presence of gametocyte exflagellation (Espinosa et al., 2013). After infection, mosquitoes were maintained in an incubator at 19-20°C and supplied with a sterile cotton pad soaked in 10% sucrose, changed every 48 hrs. SPZ were harvested 20-22 days after blood feeding.

## METHOD DETAILS

### Production of recombinant PfCSP protein constructs

Recombinant PfCSP constructs were produced as previously described (Wang et al., 2020). Briefly, the three rPfCSP constructs (FL, 5/3, Δ(NVDP)_4_) used in this study were cloned into the same CMV/R-expression vectors with a C-terminal AviTag, HRV3C-processing tag, and a 6X histidine tag (GenScript). rPfCSP constructs were expressed through transient transfection in 293F cells using the Freestyle 293F expression system (Thermo Fisher Scientific) at 37°C, 8% CO_2_ for 6 days, and purified from culture supernatants through polyhistidine-tag affinity chromatography followed by size exclusion chromatography (SEC) on an ÄKTA_TM_ Start (GE Healthcare). Monomer-containing fractions were pooled, concentrated, snap frozen, and stored at −80°C.

### Production of PfCSP peptides

All peptides used in this study were produced by direct synthesis and biotinylated by GenScript. These include 15mer peptides numbered 20-61 that were 15 amino acids in length and overlapped by 4 residues spanning the central repeat region of PfCSP, a 36mer peptide (NANP)_9_, a 31mer junctional peptide 21-25 (NPDPNANPNVDPNANPNVDPNANPNVDPNAN), 16mer peptides (NPNV)_4_ and (NPNA)_4_, and an 8mer peptide NANPNVDP.

### Isolation of plasmablasts

Plasmablasts were isolated from freshly isolated peripheral blood mononuclear cells (PBMCs) collected seven days after Sanaria PfSPZ Vaccine immunization as previously described (Kisalu et al., 2018). Briefly, PBMCs were stained for viability with Aqua LIVE/DEAD (Thermo Fisher Scientific) followed by surface staining of the following markers: CD3-PE/Cy7 (BD Bioscience), CD19-FITC (BD Bioscience), CD20-APC/Cy7 (BD Bioscience), CD27-APC (Thermo Fisher Scientific), and CD38-PE (BD Bioscience). Plasmablasts were gated as live CD3^-^CD20^-^ CD19^+^CD27^+^CD38^+^ and single cell sorted using a BD FACS Aria II (BD Immunocytometry Systems) into 96-well PCR plates containing 20 μL/well of RT reaction buffer from the SuperScript™ First-Strand Synthesis System for RT-PCR (Thermo Fisher Scientific). Plates were snap frozen on dry ice and stored at -80°C.

### Production of recombinant immunoglobulins

Recombinant immunoglobulins were produced as previously described (Wang et al., 2020). Briefly, RNA from lysed plasmablasts was reverse transcribed to cDNA (SuperScript First-Strand Synthesis System; Thermo Fisher Scientific) and immunoglobulin variable regions heavy and kappa chains were amplified using primer cocktails (Wang et al., 2020), sequenced (ACGT), and cloned into human IgG1 expression vectors (GenScript). Sequence analysis was performed using The International Immunogenetics Information System (IMGT, http://www.imgt.org/), with clonality being defined as having the same V/J genes, HCDR3 length, and HCDR3 sequence with >80% identity. Matched heavy and light chain constructs were co-transfected into Expi293 cells using the ExpiFectamine^TM^ 293 Transfection Kit (Thermo Fisher Scientific) and cultures were incubated at 37°C, 8% CO_2_ for 6 days. Supernatants were harvested and IgG was purified using rProtein A Sepharose Fast Flow resin (GE Healthcare) and buffer exchanged with 1X PBS (pH 7.4) before being concentrated using Amicon Centrifugal Filters (Millipore). Purified IgG concentrations were determined using a Nanodrop spectrophotometer.

### Determination of L9_MRCA_ sequence

The heavy chain and kappa chain sequence of L9_MRCA_ were determined using a previously described informatics pipeline for antibodyome analysis (Zhu et al., 2013). The variable (V), diversity (D), joining (J), and V(D)J gene recombination were assigned and analyzed by IgBLAST (Ye et al., 2013). Multiple sequence alignments of the germline gene and sequences related to L9 were aligned by mafft (Katoh and Standley, 2013) and the MRCA was computed using dnaml with default parameters (Felsenstein, 1989).

### Fab production and size exclusion chromatography

Purified recombinant IgGs were mixed with LysC (NEB) at a ratio of 1 μg LysC per 10 mg of IgG and incubated at 37°C for 18 hrs with nutation. The cleaved product was incubated with Protein A resin (GoldBio) at a ratio of 1 mL resin per 10 mg of initial IgG and incubated at room temperature for 1 hr to bind any uncleaved IgG and digested Fc. The purified Fab was further purified by SEC using a HiLoad 16/600 Superdex 200 pg column (GE Healthcare). Purified Fabs were incubated with 2-fold molar excess of peptide 22 and incubated at room temp for 1 hr. Fab-peptide complexes were run on a HiLoad 16/600 Superdex 200 pg column (GE Healthcare).

### ELISA for binding of mAbs to rPfCSP

Immulon 4HBX flat bottom microtiter plates (Thermo Fisher Scientific) were coated with 100 μL per well of rPfCSP to achieve molar equivalency (FL, 0.5 μg/mL; 5/3, 0.337 µg/mL; Δ(NVDP)_4_, 0.499 µg/mL) in bicarbonate buffer overnight at 4°C. Coated plates were blocked with 200 μL of PBS + 10% FBS for 2 hrs at room temp, followed by incubation for 2 hrs at 37°C with 100 μL of PfCSP or control mAbs at varying concentrations (5x10^-7^ – 5.0 μg/mL, 10-fold serial dilutions in PBS + 10% FBS). Plates were incubated with 100 μL/well of 0.1 μg/mL HRP-conjugated goat anti–human IgG (Bethyl Laboratories). Plates were washed six times with PBS + 0.05% Tween-20 between each step. After a final wash, samples were incubated for 10 min with 1-Step Ultra TMB-ELISA Substrate (Thermo Fisher Scientific). The optical density was read at 450 nm after addition of stopping solution (2N sulfuric acid, 100 μL/well).

### ELISA for binding of mAbs to peptide 22 and (NANP)_9_

Pierce™ Streptavidin Coated High Capacity Plates (Thermo Fisher Scientific) were coated with 100 μL per well of peptides (0.01 µg/mL) diluted in wash buffer (Tris-buffered saline [25mM Tris, 150mM NaCl; pH 7.2], 0.1% BSA, 0.05% Tween-20) for 2 hrs at room temp, followed by incubation for 2 hrs at 37°C with 100 μL of PfCSP or control mAbs at varying concentrations (5x10^-7^ – 5.0 μg/mL, 10-fold serial dilutions in wash buffer). Plates were incubated with 100 μL/well of 0.1 μg/mL HRP-conjugated goat anti–human IgG (Bethyl Laboratories) in wash buffer. Plates were washed three times with 200 µL wash buffer between each step. After a final wash, samples were incubated for 10 min with 1-Step Ultra TMB-ELISA Substrate (Thermo Fisher Scientific). The optical density was read at 450 nm after addition of stopping solution (2N sulfuric acid, 100 μL/well).

### Competitive ELISA

For competition of binding to rPfCSP (FL, 5/3, Δ(NVDP)_4_) by junctional peptide 21-25 and (NANP)_9_, Immulon 4HBX flat bottom microtiter plates (Thermo Fisher Scientific) were coated with rPfCSP (FL, 5/3, or Δ(NVDP)_4_; 9 nM) for 1 hr at 37°C and then blocked with 5% BSA. PfCSP mAbs were preincubated for 2 hrs at 37°C with varying concentrations (0 – 1,000 μg/mL) of either the junctional peptide 21-25 or the (NANP)_9_ peptide before being added to the coated plates. Plates were then incubated for 1 hr at room temperature with 1:20,000 dilution of HRP-conjugated goat anti–human IgG (Thermo Fisher Scientific). The plates were washed five times with PBS-Tween between each step. After a final wash, samples were incubated for 10 min with 1-Step Ultra TMB-ELISA Substrate (Thermo Fisher Scientific). The optical density was read at 450 nm after addition of stopping solution (2N sulfuric acid). Competition of binding to rPfCSP_FL_ by peptides 20-61, (NPNA)_4_ and (NPNV)_4_ was performed as described above, with plates coated with 200 ng/mL of rPfCSP_FL_ and peptide concentrations ranging from 0 – 1,000 μg/mL.

### Alanine scan with peptide 22 variants

Alanine scanning mutagenesis competitive ELISA was performed as previously described (Wang et al., 2020) using the Meso Scale Discovery (MSD) platform. Briefly, Streptavidin Multi Array Plates (Meso Scale Discovery) were coated with 200 ng/mL of rPfCSP_FL_ for 1 hr at room temperature. After coating, PfCSP mAbs preincubated for 2 hrs at 37°C with varying concentrations (0 – 1,000 μg/mL) of PfCSP peptide 22 variants where each residue was mutated to an alanine or a serine if the original residue was an alanine (GenScript) were added to the coated plates for 1 hr at room temperature. Plates were then incubated with Sulfo-tag goat anti– human IgG (Meso Scale Discovery) for 1 h at room temperature. Plates were washed five times with PBS-Tween between each step. After the final wash, plates were analyzed on an MSD Sector Image 6000 instrument after the addition of 1X MSD Read T Buffer (Meso Scale Discovery).

### Biolayer interferometry kinetic binding assay

Antibody binding kinetics were measured using biolayer interferometry (BLI) on an Octet HTX instrument (FortéBio) using streptavidin-capture biosensors (fortéBio). PfCSP mAb solutions were plated in black tilted-bottom 384-well microplates (fortéBio); assays were performed with agitation at 30°C. mAb serial concentrations used are as follow: 1.25, 0.625, 0.3125, and 0.15625 µg/mL. Loading of biotinylated peptide 22 was performed for 300 sec, followed by dipping of biosensors into buffer (PBS + 1% BSA) for 60 sec to assess baseline assay drift. Association with whole IgG (serially diluted from 16.67 to 1.04 μM) was done for 300 sec, followed by a dissociation step in buffer for 600 sec. Background subtraction of nonspecific binding was performed through measurement of association in buffer alone. Data analysis and curve fitting were performed using Octet software, version 7.0. Experimental data were fitted with the binding equations describing a 1:1 analyte-ligand interaction. Global analyses of the complete data sets, assuming binding was reversible (full dissociation), were carried out using nonlinear least-squares fitting allowing a single set of binding parameters to be obtained simultaneously for all concentrations of a given mAb dilution series.

### Isothermal titration calorimetry

Isothermal titration calorimetry was carried out using a VP-ITC microcalorimeter (Malvern Panalytical). In all titration experiments, the peptides, rPfCSP constructs and mAbs were prepared in 1X PBS, pH 7.4. Each antibody solution, prepared at a concentration of ∼40 µM (expressed per antigen binding site), was injected in 5 or 7 µl aliquots into the calorimetric cell containing the respective rPfCSP construct or peptide. The concentration of rPfCSP was ∼ 0.4 µM for the respective FL and Δ(NVDP)_4_ constructs and ∼0.8 µM for the 5/3 construct. The concentration of either of the peptides 22, 25, 29 and NANPNVDP was ∼2.5 µM. All titrations were performed at 25°C. The exact concentrations of the reactants in each experiment were determined from the absorbance at 280 nm. The heat evolved upon each injection of antibody was obtained from the integral of the calorimetric signal. The heat associated with binding to the different rPfCSP constructs was obtained by subtracting the heat of dilution from the heat of reaction. The individual heats were plotted against the molar ratio, and the enthalpy change, *ΔH*, the association constant, *K_a_* (the dissociation constant, *K_d_* =1/*K_a_*) and the stoichiometry (valency of antigen binding sites), *N*, were obtained by nonlinear regression of the data to a model that takes into account the binding to either one or two sets of sites with different binding affinities. Gibbs energy, *ΔG*, was calculated from the relation *ΔG* = -*RT*ln*K_a_*, where *R* is the universal gas constant, (1.987 cal/(K × mol)) and *T* the absolute temperature in Kelvin. The entropy contribution to Gibbs energy, *-TΔS*, was calculated from the known relation *ΔG* = *ΔH* - *TΔS.* The results were expressed per mole of antigen binding sites and the stoichiometry, *N*, denotes the number of antigen binding sites per mole of the respective rPfCSP construct.

### Crystal screening and structure determination

Purified Fabs were incubated with 1.5 molar excess of the synthetic NANPNVDP peptide (Genscript). Initial crystal screening was performed by sitting-drop vapor-diffusion in the MCSG Crystallization Suite (Anatrace) and ProPlex (Molecular Dimensions) using a NT8 drop setter (Formulatrix). For apo-L9, crystals grew in ProPlex E5 and diffracting crystals were harvested directly from screen and cryoprotected using Paratone-N. Data was collected at Advanced Light Source (ALS) beamline 5.0.2 at 12398keV. For F10_H_L9_K_-NANPNVDP, diffracting crystals were grown in a mother liquor (ML) containing 0.1M sodium citrate, pH 3.5, 1.5M (NH_4_)_2_SO_4_. Crystals were cryoprotected in ML supplemented with 30% ethylene glycol. Diffraction data was collected at ALS beamline 5.0.2 at 12286keV. For L9_H_F10_K_-NANPNVDP, diffracting crystals were grown in a ML containing 0.1M MES, pH 6.5, 18% PEG 4K, and 0.6M NaCl. Crystals were cryoprotected in ML supplemented with 30% ethylene glycol. Diffraction data was collected at Advanced Photon Source beamline 19-ID at 12669keV. All datasets were processed using XDS (Kabsch, 2010) and data reduction was performed using AIMLESS in CCP4 (Winn et al., 2011) to a resolution of 2.98Å, 1.89Å, and 2.23Å for apo-L9, F10_H_L9_K_-NANPNVDP, and L9_H_F10_K_-NANPNVDP, respectively. Initial phases for apo-L9 were solved by molecular replacement (MR) using Phaser in Phenix (Liebschner et al., 2019) with a search model of 1210 Fab (PDBid: 6D01) divided into Fv and Fc portions. The model of L9 was used as the search model for the MR of the chimera structures using Phaser. Model building was completed using Coot (Emsley et al., 2010) and refinement was performed in Phenix. The data collection and refinement statistics are summarized in Table S2. Structural figures were made in PyMOL (Schrodinger, LLC).

### Measurement of PfCSP mAb binding to SPZ

Salivary glands containing SPZ were dissected as previously described (Flores-Garcia et al., 2019) into PBS containing the protease inhibitor E-64 (Millipore-Sigma). Freshly harvested SPZ were stained with SYBR Green (10,000X concentrate; Thermo Fisher Scientific) diluted 1:2,000 in PBS for 30 min at 4°C, washed twice, and ∼8,000 SPZ were aliquoted to each well of a 96-well V-bottom plate (50 µl/well). SPZ were incubated for 30 min at 4°C with PfCSP mAbs or isotype control in PBS + 10% FBS (PBS-FBS), washed twice with 200 μl PBS-FBS, stained for 20 minutes at 4°C with goat anti-human IgG-Alexa Fluor® 647 secondary antibody (Thermo Fisher Scientific) diluted 1:1,000 in PBS-FBS, washed once with 200 μl PBS-FBS, and fixed in 200 µL PBS + 0.5% paraformaldehyde. Following fixation, events were acquired on a modified LSR II (BD Biosciences) and analyzed using FlowJo v.10.

### IV challenge with Pb-PfCSP-SPZ

To measure mAb neutralization of SPZ and reduction of parasite liver burden *in vivo*, 100 µg of PfCSP mAbs diluted in sterile filtered 1X PBS (pH 7.4; total volume 200 μL/mouse) were injected into the tail veins of female 6- to 8-week old B6(Cg)-Tyrc-2J/J albino mice (The Jackson Laboratory). After 2 hours, mice were intravenously challenged in the tail vein with 2,000 freshly harvested Pb-PfCSP-GFP/Luc-SPZ (Flores-Garcia et al., 2019). 40-42 hours post-challenge, mice were injected intraperitoneally with 150 μL of D-Luciferin (30 mg/mL), anesthetized with isoflurane, and imaged with the IVIS® Spectrum *in vivo* imaging system (PerkinElmer) 10 minutes after luciferin injection. Liver burden was quantified by analyzing a region of interest (ROI) in the upper abdominal region and determining the total flux or bioluminescent radiance (photons/sec) expressed by Pb-PfCSP-GFP/Luc-SPZ using the manufacturer’s software (Living Image 4.5, PerkinElmer).

### QUANTIFICATION AND STATISTICAL ANALYSIS

Unless otherwise indicated, all data were plotted using GraphPad Prism, version 7.0. Statistical tests used, exact value of n, what n represents, and precision measures can be found in figure legends. Unless otherwise stated, all mAbs were compared for significance to untreated/isotype control or to each other using the Kruskal-Wallis test with Dunn’s post-hoc correction for multiple comparisons or the two-tailed Mann-Whitney test for single comparisons. For the ITC stoichiometry data, errors with 95% confidence were estimated from the fits of the data. Statistics for X-ray crystal structures were calculated by Phenix and MolProbity and are summarized in Table S2.

## SUPPLEMENTAL INFORMATION

**Figure S1.**
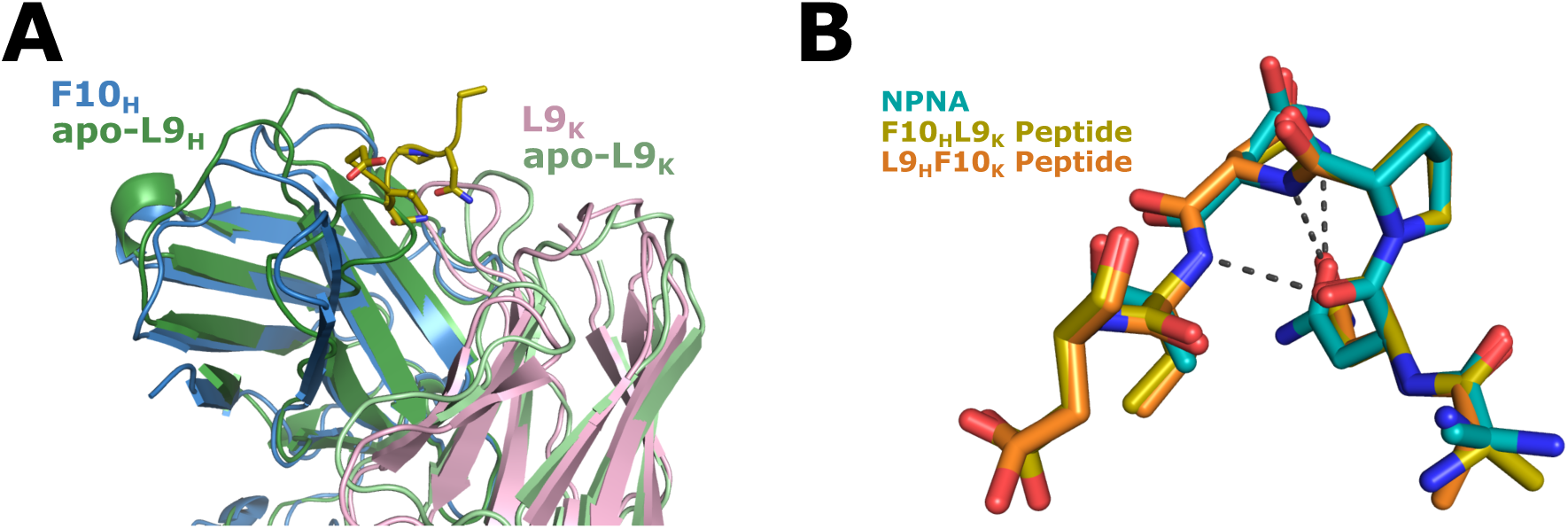
Structure of apo-L9 Fab and comparison of NANPNVDP peptide to NPNA type- 1 β-turn, related to Figure 2. (A) The structure of apo-L9 Fab aligned to F10_H_L9_K_ Fab. (B) Alignment of NANPNVDP peptide to crystal structure of NPNA, overall RMSD of 0.089 Å^2^ over four Cα atoms, showing the near-identical type-1 β-turn.

**Figures S2.**
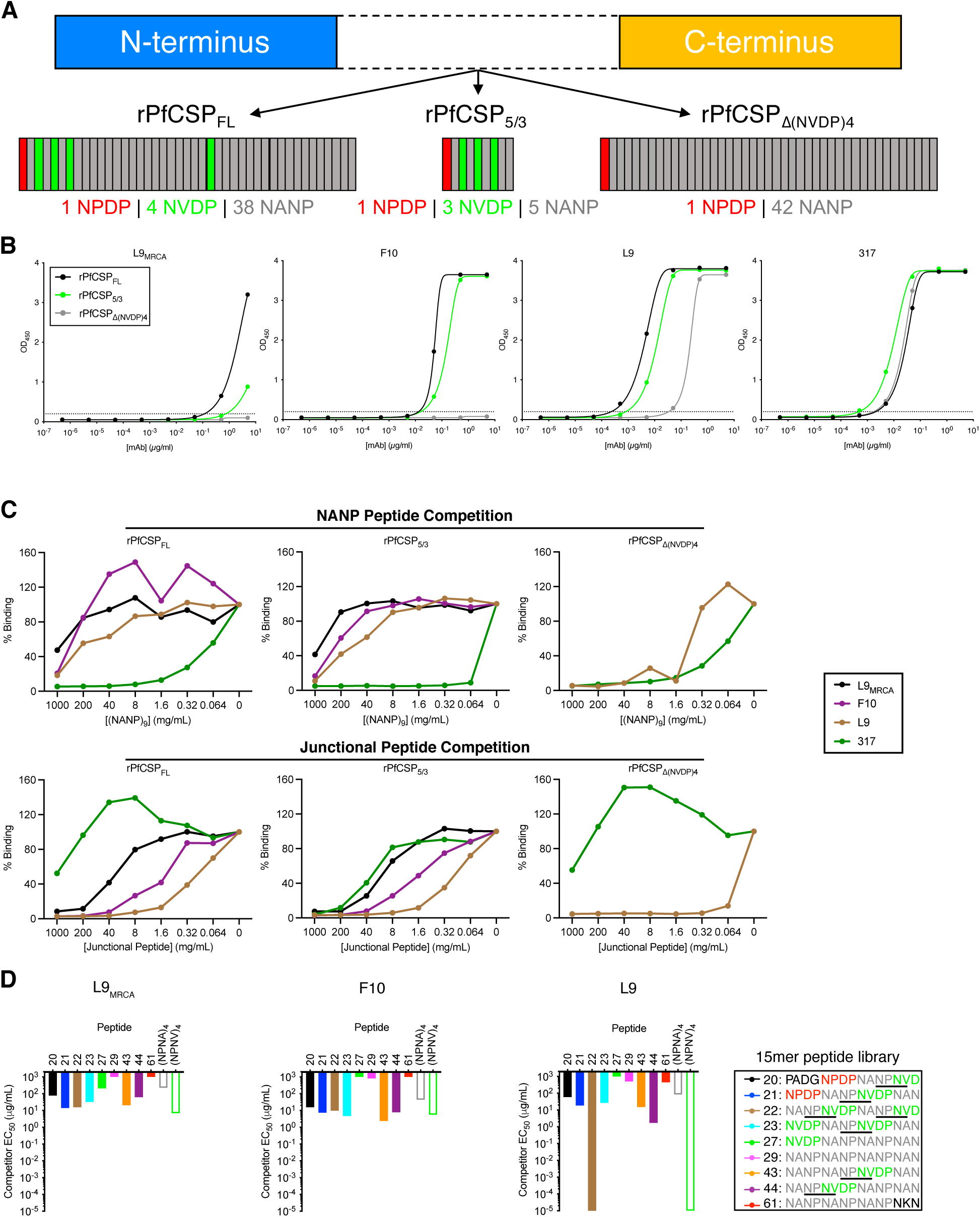
The L9 B-cell lineage has higher baseline affinity for the junctional region than for NANP repeats, related to Figure 1 and Table 1. (A) Schematics of rPfCSP constructs with identical N- and C-termini but different repeat regions (rPfCSP_FL_: 1 NPDP, 4 NVDP, 38 NANP; rPfCSP_5/3_: 1 NPDP, 5 NANP, 3 NVDP; rPfCSP_Δ(NVDP)4_: 1 NPDP, 42 NANP). (B) Binding of various concentrations of indicated mAbs to rPfCSP_FL,_ rPfCSP_5/3_, and rPfCSP_Δ(NVDP)4_ determined through ELISA; OD_450_ is plotted. (C) Competition ELISA of indicated mAbs binding to rPfCSP_FL,_ rPfCSP_5/3_, and rPfCSP_Δ(NVDP)4_ in the presence of varying concentrations of junctional peptide 21-25 (NPDPNANPNVDPNANPNVDPNANPNVDPNAN) and (NANP)_9_ peptide; OD_450_ is plotted. (D) Competition ELISA of indicated mAbs binding to rPfCSP_FL_ in the presence of varying concentrations of 15mer color-coded peptides numbered 20-61 and 16mer peptides (NPNA)_4_ and (NPNV)_4_. The half-maximal effective concentration (EC_50_) for competition by each peptide is depicted.

**Table S1, related to Figure 1.**
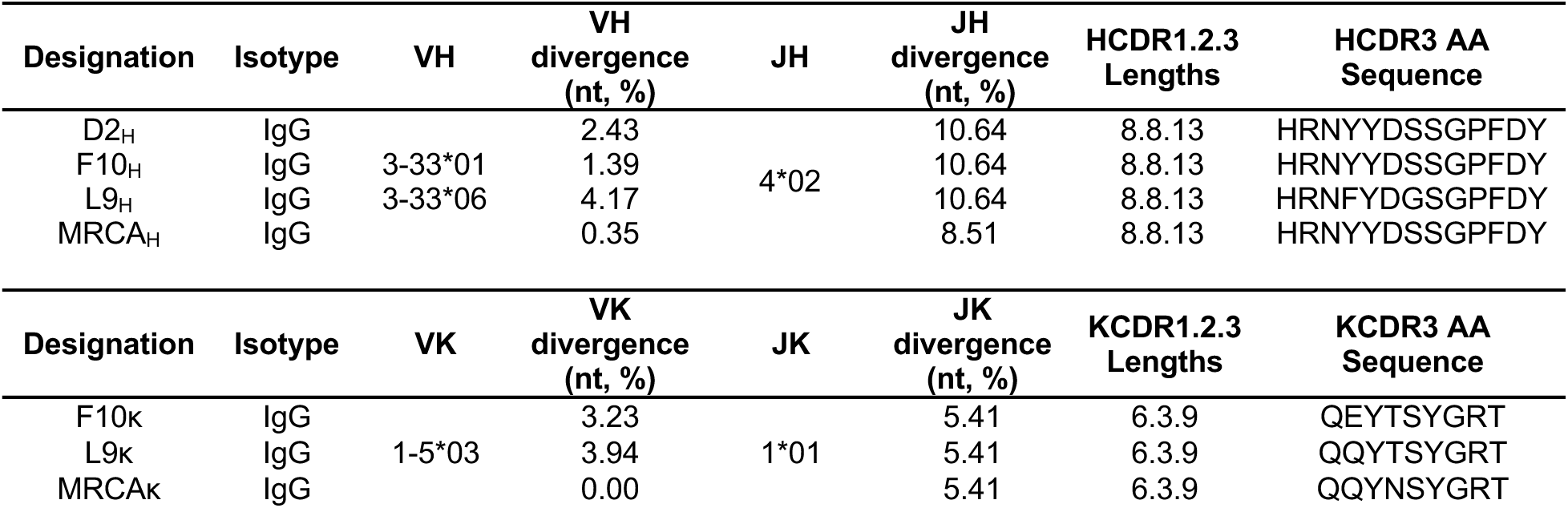
Immunoglobulin V/J-gene family usage of L9 lineage mAbs. mAb designation, isotype, V_H_/Vκ and percent divergence from germline, J_H_/Jκ gene families percent divergence from germline, V_H_/Vκ CDR1.2.3 lengths, and HCDR3/KCDR3 amino acid sequences of the L9 clonal relatives isolated in this study. n.t. = nucleotide, H = heavy, κ = kappa, J = joining, AA = amino acid.

**Table S2, related to Figures 2 and S2.** Data collection and refinement statistics for crystal structures. ^a^ R_merge_ = [∑_h_∑_i_|*I*_h_ – *I*_hi_|/∑_h_∑_i_*I*_hi_] where *I*_h_ is the mean of *I*_hi_ observations of reflection *h*. Numbers in parenthesis represent highest resolution shell. ^b^ R_factor_ and ^c^ R_free_ = ∑||F_obs_| - |F_calc_|| /∑|F_obs_| x 100 for 95% of recorded data (R_factor_) or 5% data (R_free_). ^c^ MolProbity reference

|  | apo-L9 | F10<sub>H</sub>L9<sub>K</sub>-NANPNVDP | L9<sub>H</sub>F10<sub>K</sub>-NANPNVDP |
| --- | --- | --- | --- |
| **X-ray Source** | ALS 5.0.2 | ALS 5.0.2 | APS 19-ID |
| **Data collection** |  |  |  |
| Space group | I222 | P12<sub>1</sub>1 | P2<sub>1</sub>2<sub>1</sub>2<sub>1</sub> |
| Cell dimensions |  |  |  |
| *a*, *b*, *c* (Å) | 67.964, 117.67, 143.665 | 42.787, 84.944, 64.243 | 71.503, 85.817, 145.456 |
| $\alpha$, $\beta$, $\gamma$ (°) | 90, 90, 90 | 90, 103.23, 90 | 90, 90, 90 |
| Resolution (Å) | 35.92-2.98 (3.087-2.98) | 41.65-1.89 (1.958-1.89) | 42.91-2.23 (2.31-2.23) |
| *R*<sub>merge</sub><sup>a</sup> | 0.018 (0.267) | 0.022 (0.202) | 0.042 (0.347) |
| <\|*I*/σ(*I*)> | 18.27 (2.35) | 12.92 (2.64) | 9.36 (2.28) |
| CC<sub>1/2</sub> | 0.999 (0.831) | 0.999 (0.908) | 0.998 (0.711) |
| Completeness (%) | 99.32 (100) | 93.60 (67.83) | 99.49 (97.82) |
| Redundancy | 2.0 (2.0) | 1.9 (1.7) | 2.0 (2.0) |
| **Refinement** |  |  |  |
| Resolution (Å) | 35.92-2.98 (3.10-2.98) | 41.65-1.89 (1.91-1.89) | 42.91-2.23 (2.28-2.23) |
| No. unique reflections | 12065 (1207) | 33535 (2404) | 44164 (4271) |
| *R*<sub>work</sub><sup>b</sup>/*R*<sub>free</sub><sup>c</sup> | 19.89/25.00<br>(30.05/38.09) | 16.11/20.40<br>(24.09/28.91) | 18.90/23.84<br>(25.40/32.56) |
| No. atoms | 3232 | 3584 | 7096 |
| Protein | 3229 | 3349 | 6658 |
| Water | 3 | 218 | 434 |
| Ligand | 0 | 17 | 4 |
| B-factors (Å<sup>2</sup>) | 113.2 | 42.18 | 40.78 |
| Protein | 113.24 | 41.85 | 40.66 |
| Water | 72.34 | 46.07 | 42.54 |
| Ligand |  | 57.28 | 55.58 |
| RMS bond length (Å) | 0.006 | 0.015 | 0.003 |
| RMS bond angle (°) | 1.05 | 1.31 | 0.57 |
| **Ramachadran Plot Statistics<sup>d</sup>** |  |  |  |
| Residues | 425 | 438 | 873 |
| Most Favored region (%) | 94.27 | 97.20 | 96.97 |
| Allowed Region (%) | 5.73 | 2.57 | 2.80 |
| Disallowed Region (%) | 0.00 | 0.23 | 0.23 |
| Clashscore | 14.42 | 2.41 | 2.44 |
| **PDB ID** | **7RQP** | **7RQQ** | **7RQR** |

